# High-efficiency discovery and structure-activity-relationship analysis of non-substrate-based covalent inhibitors of S-adenosylmethionine decarboxylase

**DOI:** 10.1101/2024.09.07.611751

**Authors:** Yuanbao Ai, Siyu Xu, Yan Zhang, Zhaoxiang Liu, Sen Liu

## Abstract

Targeted covalent inhibitors (TCIs) form covalent bonds with targets following initial non-covalent binding. The advantages of TCIs have driven a resurgence in rational TCI design over the past decade, resulting in the approval of several blockbuster covalent drugs. To support TCI discovery, various computational methods have been developed. However, accurately predicting TCI reactivity remains challenging due to interference between non-covalent scaffolds and reactive warheads, leading to inefficiencies in computational screening and high experimental costs. In this study, we enhanced the SCARdock protocol, a validated computational screening tool developed by our lab, by incorporating quantum chemistry-based warhead reactivity calculations. By integrating these calculations with non-covalent docking scores, docking ranks, and bonding-atom distances, non-covalent and covalent inhibitors of S-adenosylmethionine decarboxylase (AdoMetDC) were correctly classified. Using the optimized SCARdock, we successfully identified twelve new AdoMetDC covalent inhibitors from 17 compounds, achieving a 70.6% hit rate. From these novel inhibitors, we analyzed the contributions of non-covalent interactions and covalent bonding, enabling a structure-activity relationship (SAR) analysis for AdoMetDC covalent inhibitors, which was previously unexplored with substrate-based inhibitors. Overall, this work presents an efficient computational protocol for TCI discovery and offers new insights into AdoMetDC inhibitor design. We anticipate that this approach will stimulate TCI development by improving computational screening efficiency and reducing experimental costs.

## Introduction

Traditional drug discovery focuses on non-covalent drugs, and covalent drugs were deliberately avoided due to concerns of off-target reactivity and idiosyncratic drug-related toxicity ^1^. Therefore, it was a big surprise when a retrospective analysis revealed that among the 71 enzymes targeted by 313 marked drugs as of 2005, 25 enzymes (35%) were inhibited by irreversible drugs ^2^. A later statistic found that over forty covalent drugs had been approved for clinical use by then ^3^, although all the covalent drugs approved before 2010 were discovered by serendipity. Pioneered by these analyses, recent years have witnessed a fast resurgence of the rational and forward discovery of targeted covalent inhibitors (TCIs) ^1^. Covalent inhibitors outperform non-covalent inhibitors by several attracting advantages including higher efficacy, prolonged duration of action, less therapy-induced resistance, and potential in addressing challenging and “undruggable” targets ^3^. Currently, more and more covalent inhibitors have successfully progressed into clinical trials ^4^, and more encouragingly, over a dozen of covalent inhibitors have been approved in the last decade, including sotorasib inhibiting K-Ras-G12C, ibrutinib inhibiting BTK, osimertinib inhibiting EGFR-T790M, and nirmatrelvir inhibiting the SARS-CoV2 main protease ^5^.

Structurally, covalent inhibitors consist of two functional parts, a non-covalent scaffold and a reactive group (“warhead”). During the protein-inhibitor binding process, the non-covalent scaffold takes the dominant role in forming the initial complex to correctly position the warhead and provide the appropriate detention time for covalent bonding ^1^. Hence, it is reasonable to separately optimize the non-covalent scaffold and the reactive warhead to tune the selectivity of a covalent inhibitor, but the electronic interplay between the scaffold and the warhead may lead to vain attempts or low successful hits ^6^. To circumvent this scenario, most studies chose to mount mildly reactive warheads on different scaffolds by experience, nonetheless the experimental validation is still heavily dependent on a trial-and-error process ^7,8^. Based upon the binding mechanism of covalent ligands with targets, we previously presented a strategy for the computational screening of covalent inhibitors named as SCAR (referred as SCARdock hereafter) ^9,10^. In SCARdock, the reactive residue in the protein is virtually mutated or eliminated to alleviate the steric clashes between the ligand and the residue before a non-covalent docking process is performed. We demonstrated that SCARdock was successful in positioning the warhead group of a covalent ligand in the site suitable for covalent bond formation. SCARdock is easy to perform and has been applied to discover and repurpose new inhibitors ^11–14^. However, a caveat of the initial SCARdock protocol is that it only considers the non-covalent binding affinity ^10^. Obviously, the warhead reactivity of a covalent ligand is indispensable for predicting its binding affinity. As a result, the initial SCARdock protocol had high false-positive ratio and experimental costs. For example, only 3 out of 19 (15.8%) tested compounds were validated covalent inhibitors in our previous work ^10^.

AdoMetDC (S-adenosylmethionine decarboxylase) is a rate-limiting enzyme in the polyamine synthesis pathway decarboxylating AdoMet (S-adenosylmethionine) to produce dcAdoMet (decarboxylated AdoMet) for supplying the aminobutyl group during the synthesis of spermidine and spermine (Figure 1A). Spermidine, spermine, and their precursor putrescine are the major polyamines in mammalian cells. Polyamines are indispensable for normal cellular functions, and tumor cells usually have elevated levels of polyamines ^15–17^. The inhibition of polyamine synthesis has been proved to be an effective strategy to perturb cellular functions ^16,18–20^, and AdoMetDC is a promising drug target in treating cancer and parasite infection ^17,21^. AdoMetDC is a pyruvoyl-dependent decarboxylase synthesized as an inactive single-chain proenzyme. For activation, the Glu67-Ser68 bond is auto-cleaved, followed by the conversion of Ser68 to a pyruvoyl residue (Pyr) ^10^. During the decarboxylation reaction, Pyr68 serves as an electron acceptor (electrophile) and forms a Schiff base with the primary amine (-NH2) in the substrate ^22^. Previous AdoMetDC inhibitors are either not specific (for example, MGBG) or not efficient (for example, MDL-73811) ^23^. Therefore, based on this Schiff base mechanism, many covalent inhibitors containing a primary amine were designed, but these covalent inhibitors are mainly the derivates of the substrate AdoMet ^10^, which might cause non-specific toxicities since AdoMet also plays important roles in many other pathways such as being the principal methyl donor in the methylation of DNA ^24^. In our previous work, we have identified three non-substrate-based covalent AdoMetDC inhibitors with SCARdock ^10^. We reasoned that it is of great value to discover more non-substrate-based covalent inhibitors for investigating the structure-function relationship (SAR) of AdoMetDC covalent inhibitors.

**Figure 1.**
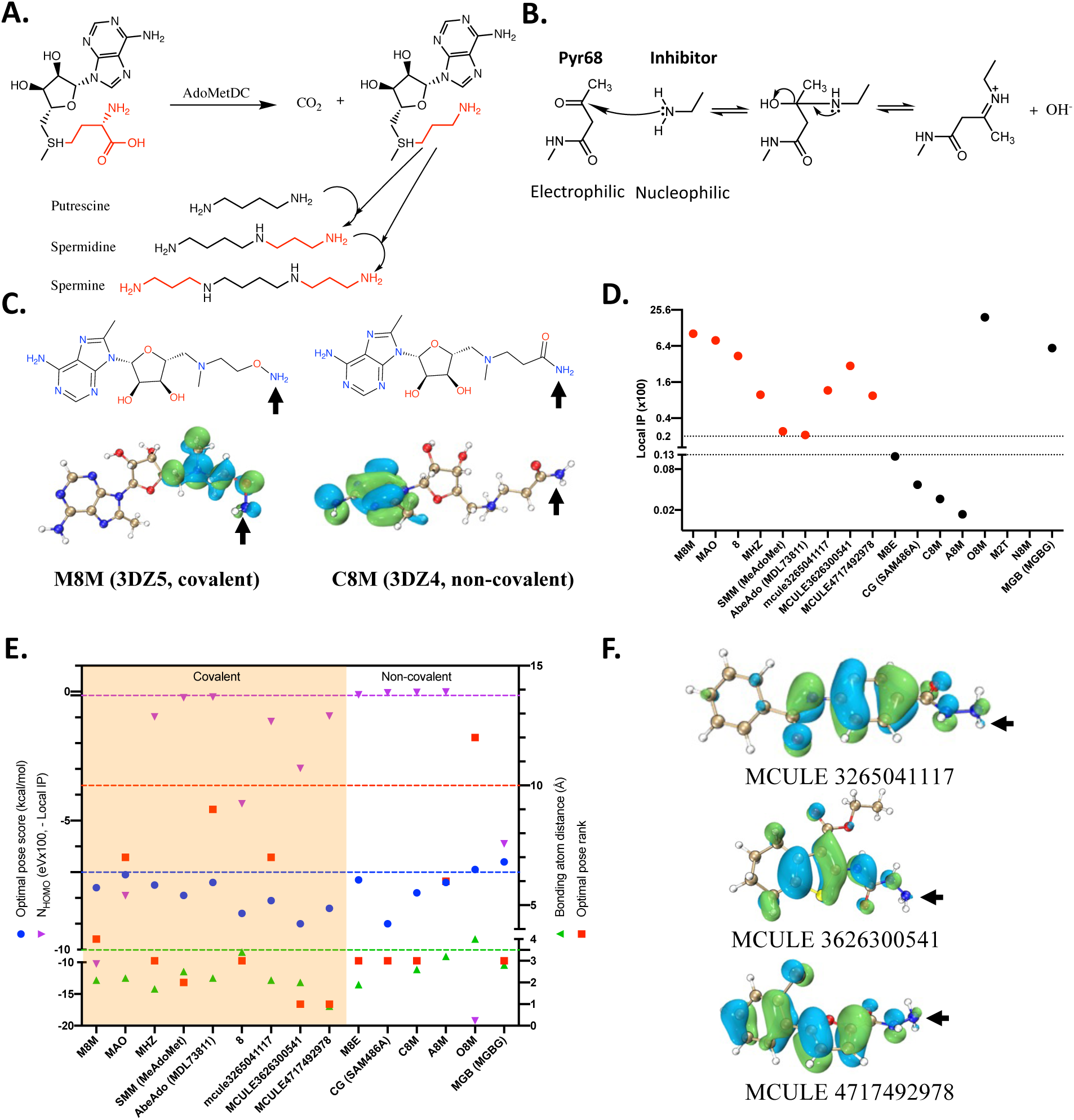
Warhead reactivity calculation helps the identification of AdoMetDC covalent inhibitors. (A) AdoMetDC catalyzes the decarboxylation of AdoMet to provide the aminopropyl group for the synthesis of spermidine and spermine. (B) The formation of the covalent bond between covalent inhibitors and Pyr68 of AdoMetDC. (C) The reactive primary amines in covalent AdoMetDC inhibitors have significant HOMO shares. Shown are two example AdoMetDC inhibitors and the other inhibitors are shown in Figure S2. The primary amines close to Pyr68 are indicated by the arrows. The HOMO orbits are shown in bubbles. The inhibitor names are shown along with the PDB IDs in the parentheses. (D) The local IP values (IP = -E_HOMO_) of the bonding atom improves the differentiation of covalent AdoMetDC inhibitors from non-covalent inhibitors. (E) The combination of four rules improves the classification of AdoMetDC inhibitors. The dotted lines are the selected thresholds for each parameter. (F) The HOMO shares of the three AdoMetDC covalent inhibitors discovered in our previous study ^10^.

In this work, we implemented a warhead reactivity prediction step in the SCARdock process. In addition, we optimized the SCARdock protocol for improved performance. Together, these optimizations significantly improved the success ratio to 70% in the discovery of non-substrate-based AdoMetDC covalent inhibitors. With these successful hits, we were able to present an SAR model of non-substrate-based AdoMetDC covalent inhibitors. Our work demonstrated the great potential of SCARdock in discovering novel covalent ligands.

## Results

### 1. Warhead reactivity calculation helps the identification of AdoMetDC covalent inhibitors

When a covalent inhibitor attacks the electrophilic pyruvoyl group (Pyr68) of AdoMetDC, the primary amine in the inhibitor contributes a lone electron pair (Figure 1B). Therefore, given a pre-defined electrophilic group (Pyr in this case), higher nucleophilicity indicates higher reactivity of the attacking ligand. It is then possible to predict the reactivity of a ligand according to its nucleophilicity. Based on molecular orbital theory and DFT (density functional theory), Jaramillo et al. ^25^ introduced an empirical (relative) nucleophilicity index definition by subtracting the HOMO (highest occupied molecular orbital) energy of a reference molecule (tetracyanoethylene) from the HOMO energy of the ligand. Since the definition of Jaramillo et al. indicates an arbitrary shifting of the origin ^25^ and the HOMO energy is a good approximation of the negative ionization potential (-IP) of a ligand ^26^, we reasoned that the IP value could be a useful descriptor for evaluating the relative nucleophilicities of a bunch of ligands. Note that this is not to re-define nucleophilicity index but to avoid the extra computation of the reference molecule. Therefore, the HOMO energies were calculated for the AdoMetDC inhibitors with experimentally verified covalent interaction. Nonetheless, the IP values (IP = - E_HOMO_) did not reliably differentiate covalent inhibitors from non-covalent inhibitors (Figure S1, Table S1). Looking into the HOMO orbitals, we noticed that the primary amines forming the Schiff base in most covalent inhibitors have significant HOMO shares (Figure 1C, Figure S2). An explanation is the local HOMO share instead of the HOMO energy of the whole molecule represents the reactivity of the bonding atom ^27^. This agrees with the reaction mechanism of the Schiff base, in which the attacking pair is the nitrogen with the lone electron pair and the positively charged carbon atom (Figure 1B). To quantitatively compare the local HOMO orbitals on the primary amines, we calculated the HOMO contribution of the nitrogen atom in the primary amine with Multiwfn ^28^. The local IP values (an indicator of the negative local HOMO energies) gave us a reasonable evaluation of the reactivity of the inhibitors. According to the local IP (referred as IP’ hereafter) values of these known inhibitors, there exists a gap (0.13-0.20) which could be a reasonable cutoff IP’ value for classifying covalent and non-covalent AdoMetDC inhibitors (Figure 1D, shown in Log_2_ scale). A note is although there were no complex structures available for AbeAdo (MDL 73811) and Compound 8, they were reported to be irreversible inhibitors of AdoMetDC ^29^. With an IP’ value cutoff between 0.13 to 0.20, the true positive (TP) value was 81.8% (9/11), the false positive (FP) value was 18.2% (2/11), the true negative (TN) value was 100% (4/4), and the false negative (FN) value was 0. The false positive prediction was from two of the non-covalent inhibitors (O8M from 3DZ7, and MGBG from 1I7C), which had IP’ values above the cutoff value.

Although a reactive warhead is necessary for covalent bonding, the warhead must be positioned appropriately by non-covalent binding. In addition, the retention time of the warhead at the bonding position must be long enough for covalent reaction. These two effectors could be respectively attributed to the distance between the bonding atoms of the warhead and the target residue, and the non-covalent docking score and ranking. Accordingly, we added three additional rules for classification: the distance between the bonding atoms of the inhibitor and the target residue (Pyr68) is no more than 3.5 Å, the SCARdock score is not higher than -7.0 kcal/mol, and the ranking of the conformation is top 10. Combining this rule with the IP’ rule, both TP and TN would be 100%, with both FP and FN being 0 (Table S1, Figure 1E). Since Compound 8 was the only positive inhibitor that had the bonding atom distance larger than 2.5 Å due to the alternative rotation of the primary amine in the optimal pose (Figure S3), we reasoned that a lower bonding atom distance threshold (2.5 Å) should be good for in silico screening. Our theoretical analysis is also a confirmation of previous hypothesis that warhead positioning and reactivity are both necessary for evaluating covalent inhibitors. To investigate whether the proposed rules is useful in the discovery of novel covalent AdoMetDC inhibitors, we calculated the IP’ values of the three covalent inhibitors discovered in our previous work^10^. Encouragingly, those three covalent inhibitors were classified correctly (Table S1, Figure 1E, 1F).

### 2. Warhead activity prediction improves SCARdock screening efficiency

Then, we asked if the combined classification rule could improve the screening efficiency of AdoMetDC covalent inhibitors using SCARdock. According to this rule, we made the following optimization to our initial protocol ^10^ during the SCARdock process (Figure 2A): (1) Pre-filter the compound libraries with the potential warheads to downsize the screening library; (2) Output only top-ten docking conformations. After docking, the results were firstly filtered by the lowest score and the score density as previously described ^10^. Next, the distance between the bonding atoms was calculated for each conformation, and the compound was kept if at least one conformation passed the threshold of the bonding atom distance (2.5 Å). Meanwhile, the docking scores of these specific conformations were evaluated to see if they were lower than the docking score threshold (-7.0 kcal/mol). To minimize the computational cost, the compounds passing these rules were sent to vendors for inquiring the availability before the molecular descriptors of the purchasable compounds were calculated. The molecules with primary amines passing the local IP’ threshold (0.2) were visually evaluated and selected. In addition, several compounds were also selected for the purpose of structure-activity relationship (SAR) analysis. In total, 18 compounds (Table S2) were purchased for experimental validation. Three compounds with slightly lower IP’ values below the threshold were included because their structures are good for structure-activity analysis later. Among these 18 compounds, one (AG-690/33251022) was not tested due to low solubility, and it was not included in future analyses.

**Figure 2.**
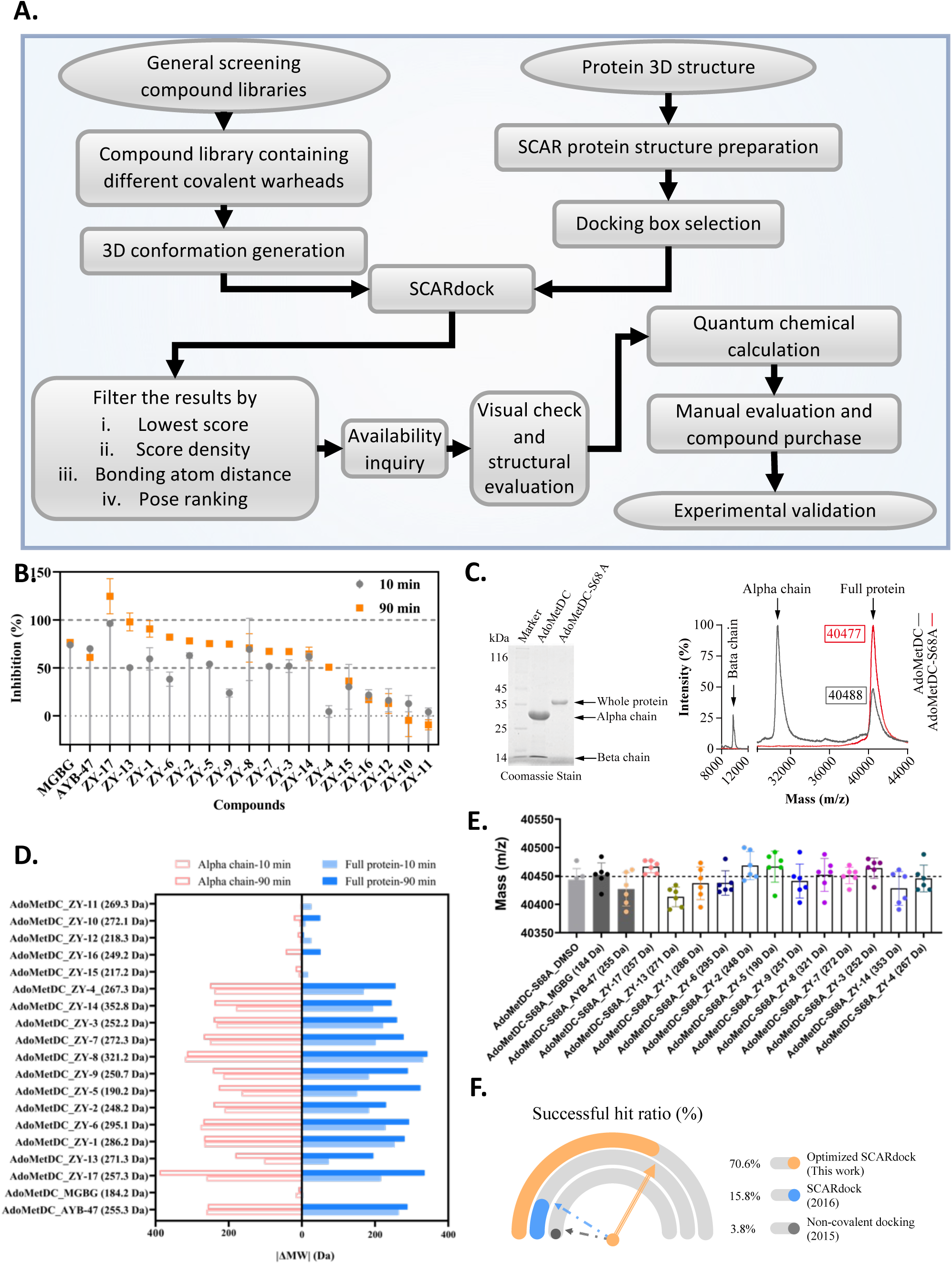
Warhead activity prediction improves SCARdock screening efficiency. (A) The diagram of the updated SCARdock process. (B) The time-dependent inhibition of the compounds on AdoMetDC activity. (C) The purified wild-type AdoMetDC has an alpha chain and a beta chain, whereas the inactive S68A mutant has a single chain as determined by SDS-PAGE and MALDI-TOF. (D) The molecular weight changes in the full protein and the alpha chain of the wild-type AdoMetDC protein as determined by MALDI-TOF. The theoretical molecular weights of the compounds are shown in the parentheses. (E) The molecular weight changes in the AdoMetDC-S68A full protein after incubation with the compounds. (F) The evolution of the successful hit ratios of our previous work ^30^ ^10^ and this work.

The inhibitory effect of these compounds was firstly evaluated by the AdoMetDC-PEPC-MDH assay we established previously ^30^. The non-covalent inhibitor MGBG and the covalent inhibitor MCULE-3265041117 (referred as AYB-47 hereafter) we identified previously were used as the controls. These 17 compounds did not significantly interfere the signal at 100 µM (Figure S4) and were named as ZY-1 to ZY-17 for simplicity (Table S2). A half of these 17 compounds (9/17, 52.9%) showed over 50% inhibitory effect on AdoMetDC activity with 10 min of compound-enzyme pre-incubation (Figure S5). When the pre-incubation time was increased to 90 min, 12 compounds (70.6%) showed over 50% inhibitory effect (Figure S6). Meanwhile, 10 compounds (58.9%) showed higher inhibitory efficiency with 90-min pre-incubation than with 10-min pre-incubation (Figure 2B).

To check if these compounds indeed form covalent complexes with AdoMetDC, we used MALDI-TOF to evaluate the molecular weights of pre-incubated samples. With 10 min of pre-incubation at the concentration ratio of 1:1 (protein vs compound), increased molecular weights were noticed for three compounds (Figure S7). When the pre-incubation concentration ratio was increased to 1:5 (10 min), 12 compounds led to increased molecular weights (Figure S8). When the pre-incubation time was extended to 90 min (1:5), no more compounds were found to cause molecular weight increases (Figure S9). Comparing the molecular weight changes in the whole protein and the separated alpha/beta chains (Figure 2C, 2D), we confirmed that 12 compounds covalently bound to the AdoMetDC alpha chain containing Pyr68 but not the beta chain (residues 1-67).

To further validate the binding site of these inhibitors, Ser68 was mutated to alanine. This mutant cannot self-activate and does not have catalytic activity. The MALDI-TOF data showed that, as expected, none of these compounds caused changes in the molecular weight of the mutant protein AdoMetDC-S68A (Figure 2E). Also, only the intact protein was noticed, confirming that AdoMetDC-S68A did not self-cleave into two sub-units.

Compare the spectrometry data and the activity data, we noticed that these two assays were consistent. Therefore, we concluded that 12 compounds can form covalent complexes with AdoMetDC, whereas 5 compounds can not. Compared to the successful hit ratio (3 in 19, or 15.8%) in our previous work, this work had a surprisingly large improvement (12 in 17, or 70.6%, excluding the compound with low solubility) (Figure 2F).

### 3. Local ionization potential is a good predictor of covalent bonding activity

We further assessed the binding between these inhibitors with AdoMetDC using the thermal-shifting assay (TSA). Both the Sypro-Orange based TSA and the gel-based TSA showed that except ZY-13, all inhibitors improved the thermal stability of AdoMetDC (Figure 3A, Figure S10, S11). The IC_50_ values of these compounds showed large variances, with ZY-13 and ZY-4 showing lowest inhibitory effects (Figure 3B, S12). To understand the contributions of non-covalent binding and covalent bonding of the covalent inhibitors, we tried to quantify the non-covalent binding step (*K*i) and the bonding step (*k*_inact_) using the AdoMetDC-PEPC-MDH assay. Limited by the temporal resolution of this assay, the kinetic data were only obtained for six inhibitors with moderate activity (Figure 3B, 3C, Figure S13). Therefore, we used the BLI assay to evaluate how non-covalent interaction contributes to the inhibitory potency of these new AdoMetDC covalent inhibitors, reasoning that a short (60 s) binding phase will majorly reflect the non-covalent binding affinity (*K*_i_) between the inhibitor and AdoMetDC. Our data (Figure 3D) showed that the non-covalent binding affinities of these inhibitors vary a lot from several millimolar to over one hundred millimolar. For the six moderate inhibitors, we were able to perform the correlation analyses (Figure 3E, 3F, Table S2), which showed good positive correlation for *K*_D_-*K*_i_ and IP’-*k*_inact_, confirming that both non-covalent binding and covalent bonding contribute to the activity of covalent inhibitors, and the IP’ value is a good indicator of the warhead activity.

**Figure 3.**
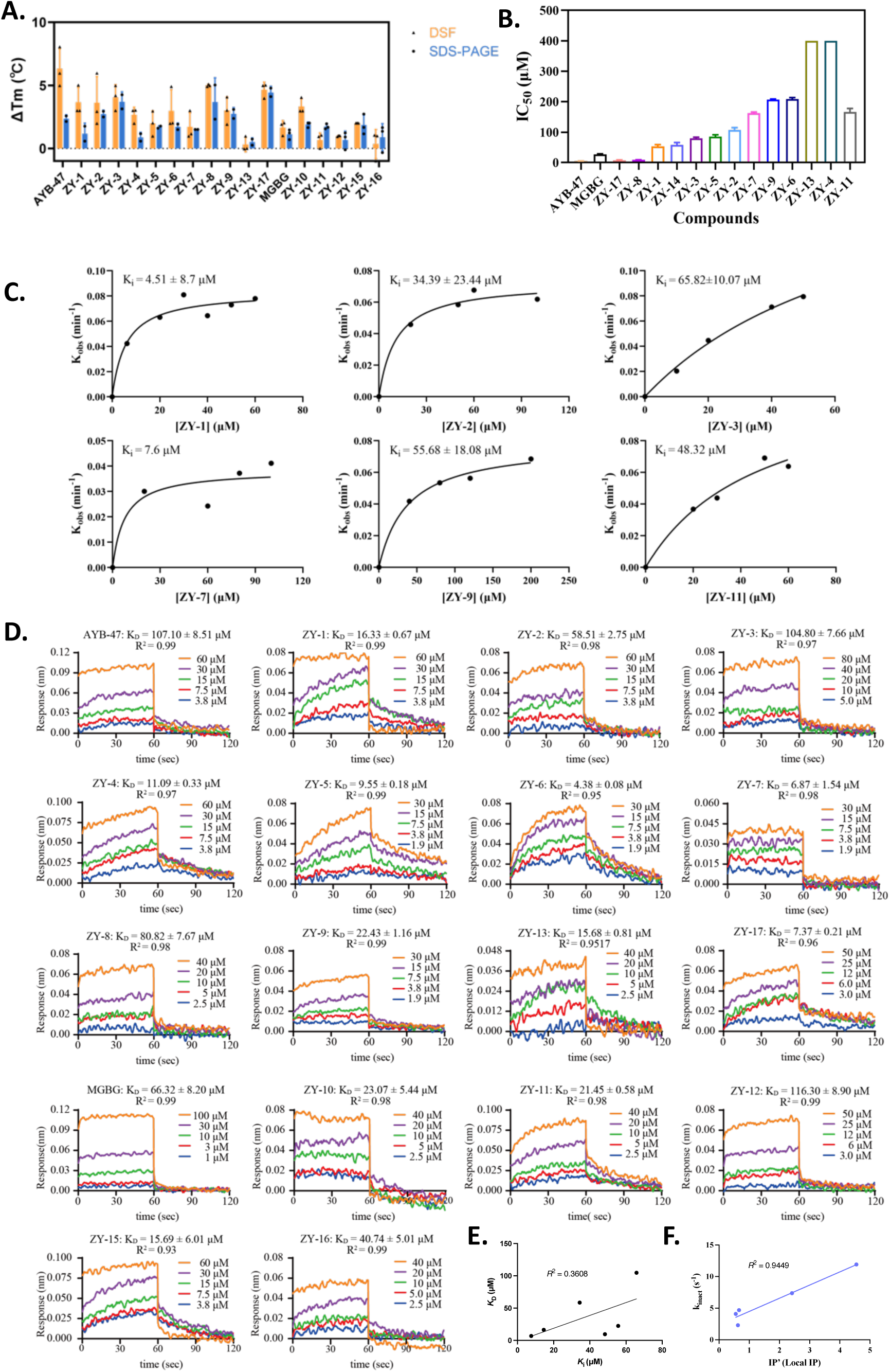
Local ionization potential is a good predictor of covalent bonding activity. (A) The binding of the compounds affects the thermal stability of AdoMetDC as evaluated by the Sypro-Orange based TSA and the gel-based TSA. (B) The IC_50_ values of the compounds were determined with the AdoMetDC-PEPC-MDH assay. The IC_50_ values of ZY-10, ZY-12, ZY-15 and ZY-16 was not obtained. (C) The kinetic data of six covalent inhibitors were determined with the AdoMetDC-PEPC-MDH assay. (D) The non-covalent binding affinities were evaluated with the BLI assay. The data for ZY-14 was not determined. (E) The *K*_D_-*K*_i_ correlation of the six moderate covalent inhibitors. (F) The IP’-*k*_inact_ correlation of the six moderate covalent inhibitors. ZY-2 was excluded because it has a very large IP’ value.

### 4. The inhibitors reveal a structure-activity relationship of non-substrate-based AdoMetDC covalent inhibitors

The data above indicated that we obtained a diverse set of non-substrate-based AdoMetDC inhibitors. Therefore, we set out to analyze their structure-activity relationship (SAR) (Figure 4A). In our screening, all covalent inhibitors have a same warhead group, because the compounds with the other warheads did not show covalent binding potency. According to the binding conformation of AdoMet, we dissect the structure of a representative AdoMetDC covalent inhibitor (MeAdoMet) into five substructures (I-V) (Figure 4B). Previous reports had analyzed the SAR of substrate-based AdoMetDC inhibitors, emphasizing the importance of the length of the substructure I and the cation substructure II. Basically, our non-substrate-based inhibitors showed similar activity elements as the substrate-based inhibitors (Figure 4C). The optimal length of I is two bonds, and the optimal length of III is three bonds. One of our non-substrate-based inhibitors, ZY-14, has a substructure V extending to a side pocket next to Pyr68. This substructure is good for improving the inhibitor’s binding affinity. A major difference between the substrate-based covalent inhibitors and our non-substrate-based covalent inhibitors is in the substructures II and IV. In the substrate-based inhibitors, II prefers a cation group, forming cation-pi interaction with F223 and/or F7. However, II is substituted by an aromatic group, which forms a pi-pi stacking interaction with F223. As of the substructure IV, in the substrate-based inhibitors, the adenosine group is bended by the interaction from the ribose to form twisted pi-pi interactions with F7. However, in our non-substrate-based inhibitors, an aromatic ring forms pi-pi stacking with F7 with a better parallel conformation (Figure 4C). Meanwhile, a trifluoromethyl group is also preferable at the substructure IV.

**Figure 4.**
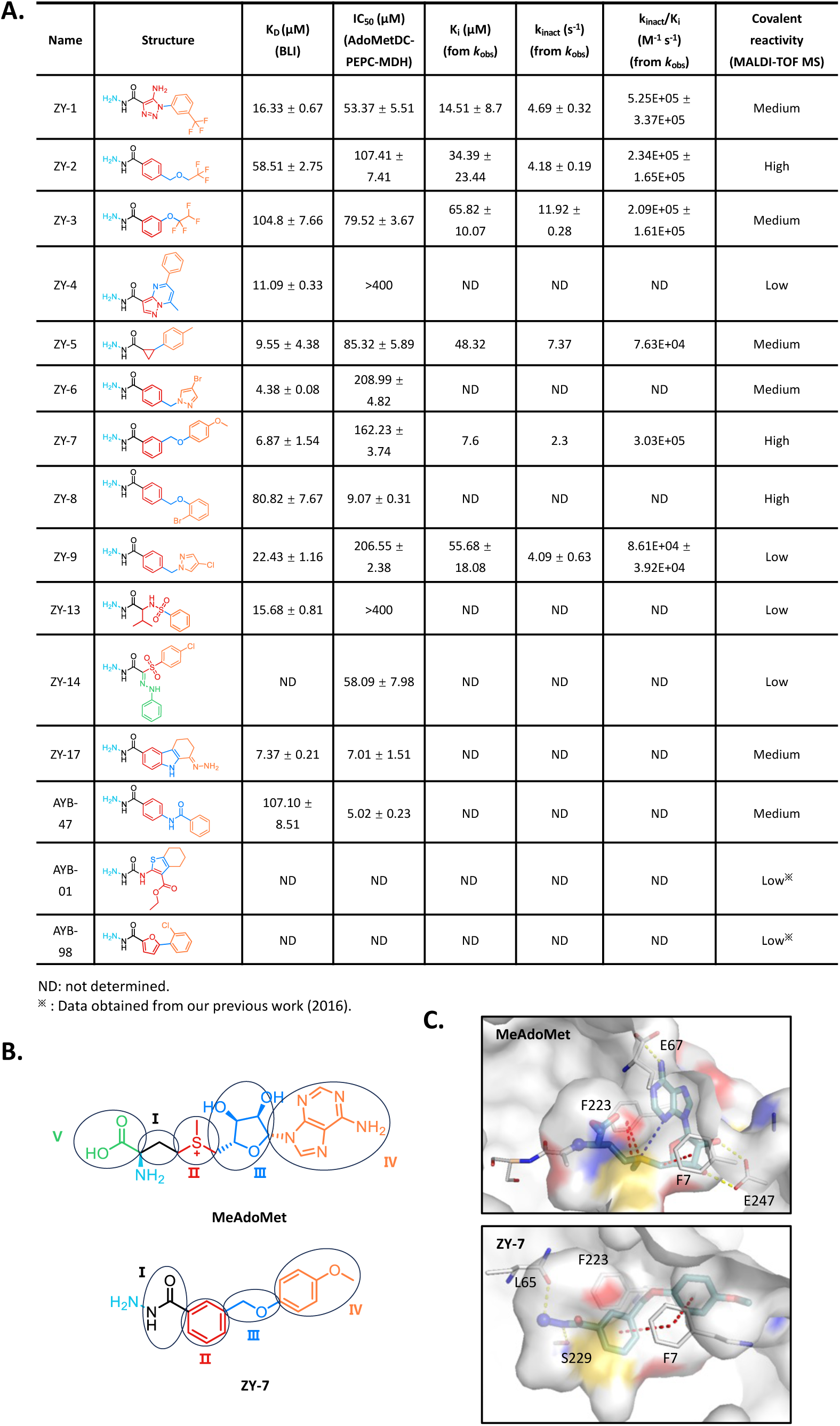
The new inhibitors reveal new structure-activity relationships. (A) The structures and the experimental data of the AdoMetDC covalent inhibitors identified in this work and our previous work^10^. (B) The comparison of the substructures of a substrate-based inhibitor (MeAdoMet) and a non-substrate-based inhibitor. The structurally equivalent substructures are in same colors. (C) The 3D binding conformations of a substrate-based inhibitor (MeAdoMet) and a non-substrate-based inhibitor. AdoMetDC is shown in surface, with key residues shown in sticks. The compounds are shown in sticks.

## Discussion

After a decade of fast development, the therapeutic advantages of targeted covalent inhibitors (TCIs) have been widely recognized ^5,31^. Compared to non-covalent inhibitors, TCIs are more complicated, because they contain a non-covalent scaffold and a reactive warhead. To affiliate the rational discovery of TCIs, many computational protocols have been developed in recent years ^32–34^. Based on the binding mechanism of TCIs, we previously presented SCARdock ^10^, a computational protocol for discovering TCIs based on well-developed non-covalent docking protocols. The efficacy of SCARdock has been experimentally validated, but like the other TCI computational protocols, its screening efficiency is not high. A major challenge for the TCI-targeted computational tools is the prediction of the bonding reactivity of TCIs. Some studies have tried to predict the reactivity of covalent inhibitors ^35–39^, but reliable methods for predicting the warhead reactivity of TCIs are still lacking. Therefore, most endeavors are still dependent on the trial-and-error protocol by experimentally testing different structural combinations of warheads and scaffolds.

In this work, we found that it is possible to classify previous AdoMetDC inhibitors into covalent inhibitors and non-covalent inhibitors by combining four rules: the distance between the bonding atoms of the inhibitor and the target residue (Pyr68), the SCARdock score, the ranking of the conformation, and the ionization potential of the bonding atom (local IP) of the inhibitor. Based on this combinatory rule, we were able to successfully identify 12 covalent inhibitors from 17 compounds, achieving a successful hit ratio of 70.6%. Since these compounds have a same warhead but different scaffolds, we were able to elucidate a structure-activity relationship of non-substrate-based AdoMetDC covalent inhibitors. We found that these compounds prefer to form pi-pi stacking interactions with AdoMetDC-F7 and AdoMetDC-F223 with two separate aromatic rings, which differs from previous substrate-based covalent inhibitors.

One caveat of this work is that the local IP values of some inhibitors did not agree with their covalent bonding activity perfectly. A major reason might be due to solvent effects, which were known to be significant in electrophile/nucleophile interactions. How to predict solvent effects would be a great challenge in the future. Meanwhile, if the local electrophilic/nucleophilic calculation could be used on the covalent inhibitors targeting the proteins with nucleophilic residues remains to be investigated further.

Taken together, we presented a protocol for discovering AdoMetDC covalent inhibitors with very high efficiency. This work, along with our previous work ^10^, confirmed that SCARdock is a valuable protocol for discovering covalent inhibitors. Future work to demonstrate that the application of SCARdock on other protein targets would be of great value to the field of covalent drug discovery.

## Author contributions

Conceptualization, S.L.; methodology, S.L.; software, S.L.; validation, S.L., Y.B.A., Y.Z, and S.Y.X.; formal analysis, S.L., Z.Y., S.Y., Z.X.L., and Y.B.A.; investigation, S.L., Z.Y., S.Y., Z.X.L., and Y.B.A.; resources, S.L.; writing-original draft preparation, S.L, S.Y.X., Y.B.A., and Y.Z.; writing-review and editing, S.L.; visualization, S.L., S.Y.X., Y.B.A., and Y.Z.; supervision, S.L.; project administration, S.L.; funding acquisition, S.L.. All authors have read and agreed to the published version of the manuscript.

## Acknowledgements

We would like to thank the support from the other members of our lab. This research was funded by the National Natural Science Foundation of China grant number [31971150], the Department of Science and Technology of the Hubei Provincial People’s Government grant numbers [2024AFA014, 2019CFA069], and the Opening Fund of Collaborative Innovation Center for Industrial Fermentation (Ministry of Education & Hubei Province) grant number [2023SL01]. We also thank Prof. Jun Gao at Huazhong Agricultural University for his help on quantum chemical calculation.

## Methods

### SCARdock2 screening

The SCARdock process was similar to our previous report ^10^ with small modifications. Briefly, the crystal structure of AdoMetDC (PDB ID: 3DZ5) was used for docking after Pyr68 was removed. The SPECS screening library (May 2020) was filtered with known warheads as described in ^9^. The 3D conformations of small molecules were generated with RDKit ^40^. Different from the original SCARdock protocol ^10^, the distance between the presumptive bonding atoms in the final form was used for distance filter.

### Quantum chemical calculation

The molecular orbitals were calculated using Gaussian16 at the density functional theory (DFT) level with the B3LYP/6-311+G(d,p) basis set. Harmonic vibrational frequencies were calculated for all stationary points. Multiwfn (v3.7) ^28^ was used for molecular orbital decomposition (SCPA method) and calculate the HOMO contribution of bonding atoms from the Gaussian data. VMD (v1.9.4) ^41^ was used for plotting the molecular orbits from the Gaussian data.

### Protein expression and purification

The human AdoMetDC was expressed and purified as previously described. The AdoMetDC S68A mutation is generated by using the FAST Site-Directed Mutagenesis Kit (Tiangen; Beijing; China).

### AdoMetDC-PEPC-MDH assay

The initial screening of AdoMetDC covalent inhibitors was carried out using the previously established AdoMetDC-PEPC-MDH assay with similar procedures. Initially, 17 commercial compounds were completely dissolved in DMSO. To validate the effect of the compounds on the PEPC-MDH assay, both the standard HCO3-and the compound concentrations were 100 µM. In the final reaction mixture of the AdoMetDC-PEPC-MDH assay, AdoMetDC was 2 μM, AdoMet was 1 mM and the compounds were 100 μM. The compounds were incubated with AdoMetDC for 10 min or 90 min at 37 °C before the reaction was initiated by transferring the mixture to the solution containing the substrate AdoMet. DMSO and MGBG (100 μM) were used as controls.

### Deriving *k*_inact_/*K*_i_ from *k*_obs_ values

The AdoMetDC-PEPC-MDH assay was used to determine the inhibition ratio of compounds at different concentrations and pre-incubation times. The *<u>k</u>*<u>_obs_</u> value was obtained with the following formula:

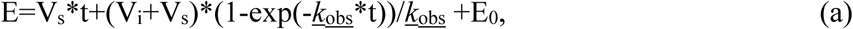

where E is the concentration of enzyme-inhibitor (*E*·*I*) complex, V_s_ is the steady generation rate of the complex, E_0_ is the concentration of the complex without pre-incubation.

Subsequently, *k*_inact_/*K*_i_ was obtained after the *k*_obs_ value was substituted into the formula below ^42,43^:

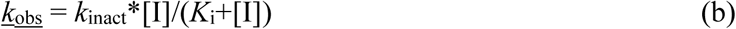

### Mass spectrometry analysis of digested proteins

The AdoMetDC protein was combined with the compounds or DMSO in a buffer solution consisting of 1 mM NaCl, 10 mM HEPES (pH 7.0) and a final reaction volume of 20 μL. MGBG and AYB-47 were used as negative and positive controls, respectively. A mixture of 20 μM AdoMetDC protein and 20 μM compounds or an equal volume of DMSO was first incubated at 37°C for 10 minutes. Subsequently, 1 μL of SA and 1 μL of the above mixture were applied to a MALDI-TOF metal plate in triplicate. In addition, the concentration of the compounds was increased to 100 μM and the incubation time was extended to either 10 or 90 min. Once the droplets had completely evaporated, the metal plates were transferred to the instrument for analysis. Flex Control software was used for data acquisition and Flex Analysis software for data processing ^44^.

### BLI (Bio-layer interferometry) assay

First, purified AdoMetDC protein was biotinylated using the Biotin Rapid Labeling Kit (Jiangsu BoMeiDa Life Science Co., LTD.) at room temperature and then diluted to 0.3 mg/mL in PBS (Hyclone), pH 7.5. The super streptavidin (SSA) biosensors were immersed in 200 μL of this solution for 300s to load the biotinylated AdoMetDC protein on their surface. The compounds were dissolved in DMSO to a concentration of 10 mM and then further diluted gradiently using detection buffer (PBS containing 0.02% Tween 20 and 1% DMSO, pH 7.5). The final concentration of DMSO was below 2%. Subsequently, the sensors were subjected to association and dissociation with solutions containing different concentrations of compounds and DMSO, respectively, for 60 s each. In addition, it is essential to establish reference samples and reference sensors to prevent non-specific binding of compounds to sensors. The above procedure was repeated at 30°C. Following the detection step, the data was processed using the ForteBio Data Analysis HT 10.0 using the double-deduction approach. Finally, KD values were calculated using steady-state analysis with 1:1 global model fitting. R^2^ > 0.95 indicated a strong linear relationship between compound concentration and response value ^45^.

### SYPRO Orange based TSA (thermal shift assay)

0.2 µL of 5000 × SYPRO Orange (Sigma-Aldrich) was mixed with AdoMetDC protein (12.5 µM final) and compounds (100 µM final) in 1 × PBS (pH = 7.5), the final volume of the mixture is 200 µL. Parallel wells of 50 µL of each solution were placed in white PCR strip tubes, which were then sealed with highly transparent, optically clear quality sealing tape (Roche). Plates were heated in a CFX96 Real-Time System (Bio-Rad) from 30 to 95 °C at a rate of 1 °C per minute. Fluorescence changes (excitation 470 nm, emission 570 nm) in each well were recorded in real time. Data were analyzed in Bio-Rad CFX Manager 3.0 software to calculate melting temperatures ^46,47^.

### SDS-PAGE based TSA

Briefly, 200 µL of a master mix containing 12.5 µM AdoMetDC protein in assay buffer (0.1 mM EDTA·2Na, 200 mM NaCl, 10 mM HEPES, 2.5 mM DTT) was treated with 2 µL of compounds (10 mM) or vehicle controls. The mixture was incubated in a CO₂ incubator at 37 ℃ for 90 minutes. After incubation, the mixture was divided into 20 µL aliquots which were then subjected to various temperatures for 3 minutes to denature the AdoMetDC proteins. After heating, the samples were centrifuged to pellet the aggregated AdoMetDC protein and the remaining soluble protein was analyzed by SDS-PAGE. AdoMetDC protein bands were quantified using ImageJ (NIH). The melting curves were analyzed using the Boltzmann sigmoid equation ^48,49^.

