## Supplementary Information for "High-efficiency discovery and structure-activity-relationship analysis of non-substrate-based covalent inhibitors of S-adenosylmethionine decarboxylase"

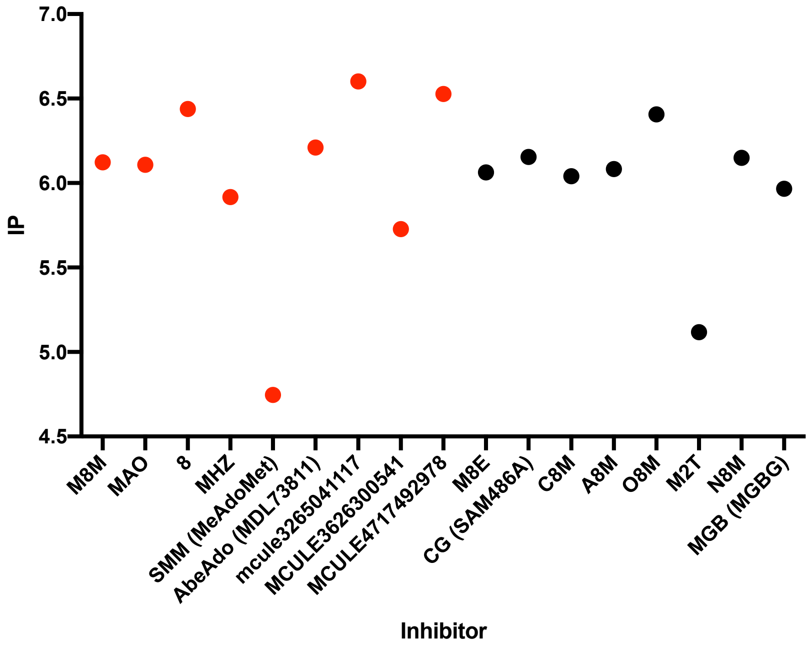


**Figure S1**. The molecular IP values (IP = - HOMO) could not differentiate covalent AdoMetDC inhibitors from non-covalent inhibitors.

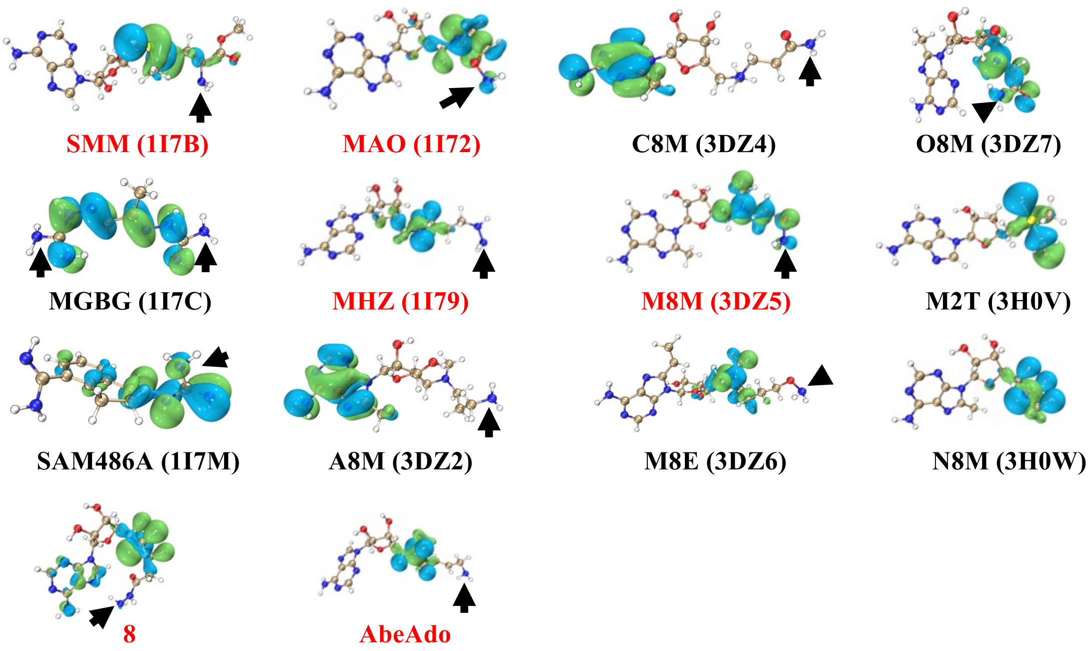


**Figure S2**. The local HOMO shares (shown in bubbles) of the known AdoMetDC inhibitors. The red names indicate covalent inhibitors. The inhibitor names are shown along with the PDB IDs in the parentheses. The primary amines close to Pyr68 are indicated by the arrows.

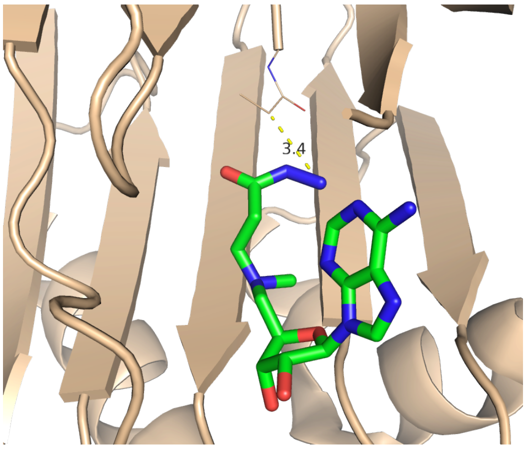


**Figure S3**. Compound 8 has a distance over 2.5 Å between the two bonding atoms from Pyr68 and the primary amine in the SCARdock result.

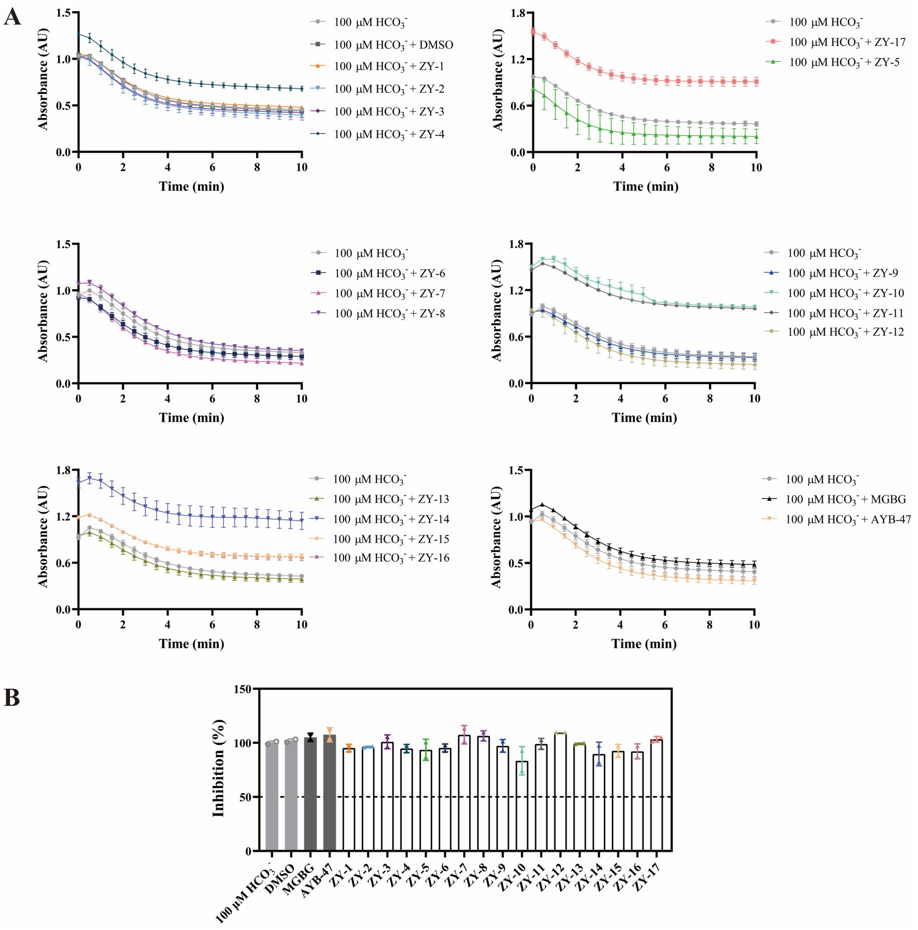


**Figure S4.** (A) The enzymatic activity of AdoMetDC was evaluated with the AdoMetDC-PEPC-MDH assay and the absorbance value of the AdoMetDC protein was monitored in real time at a wavelength of 340 nm with the standard $HCO_3^-$ and the compounds. (B) The inhibitory efficiency of compounds on AdoMetDC activity.

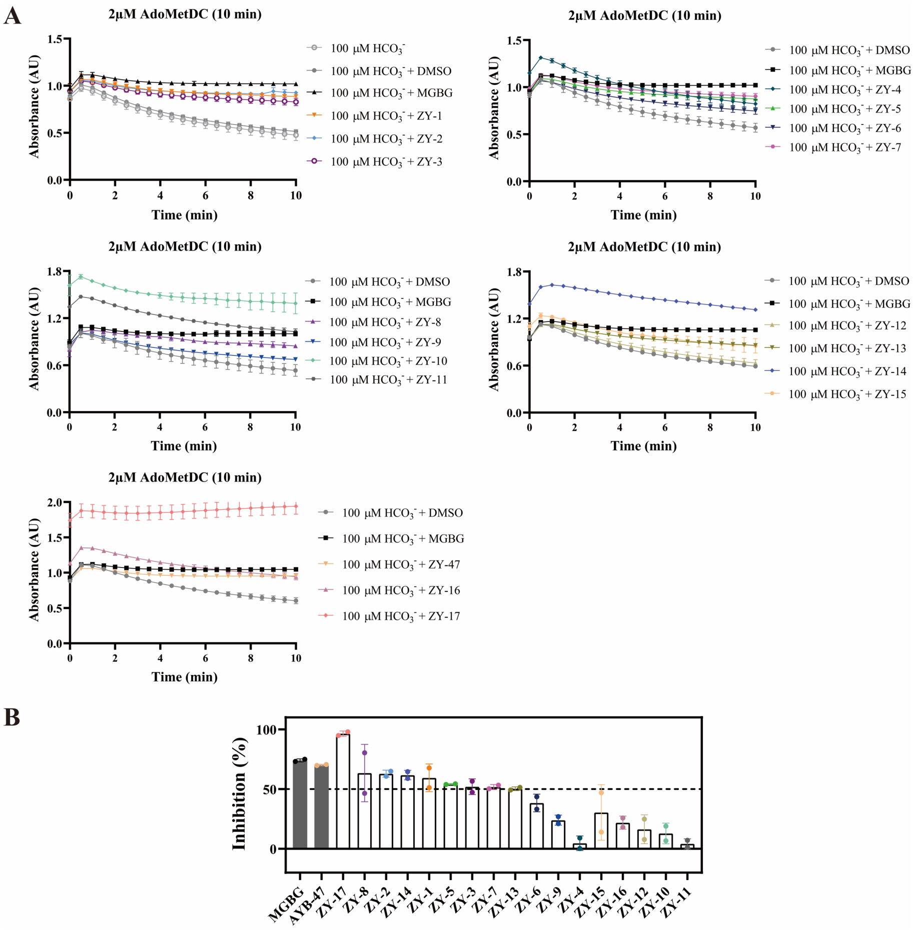


**Figure S5.** (A) The activity of AdoMetDC was monitored in real time at a wavelength of 340 nm with 10 min of pre-incubation with the compounds. (B) The inhibitory efficiency of compounds with 10-min pre-incubation.

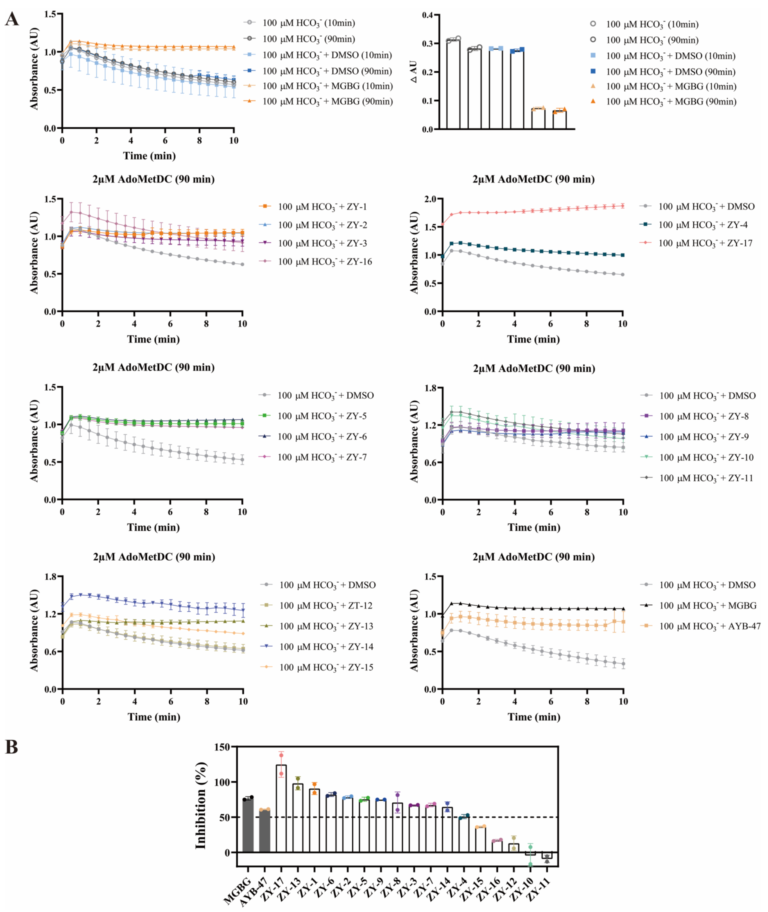


**Figure S6.** (A) The activity of AdoMetDC was monitored in real time at a wavelength of 340 nm with 90 min of pre-incubation with the compounds. (B) The inhibitory efficiency of compounds with 90-min pre-incubation.

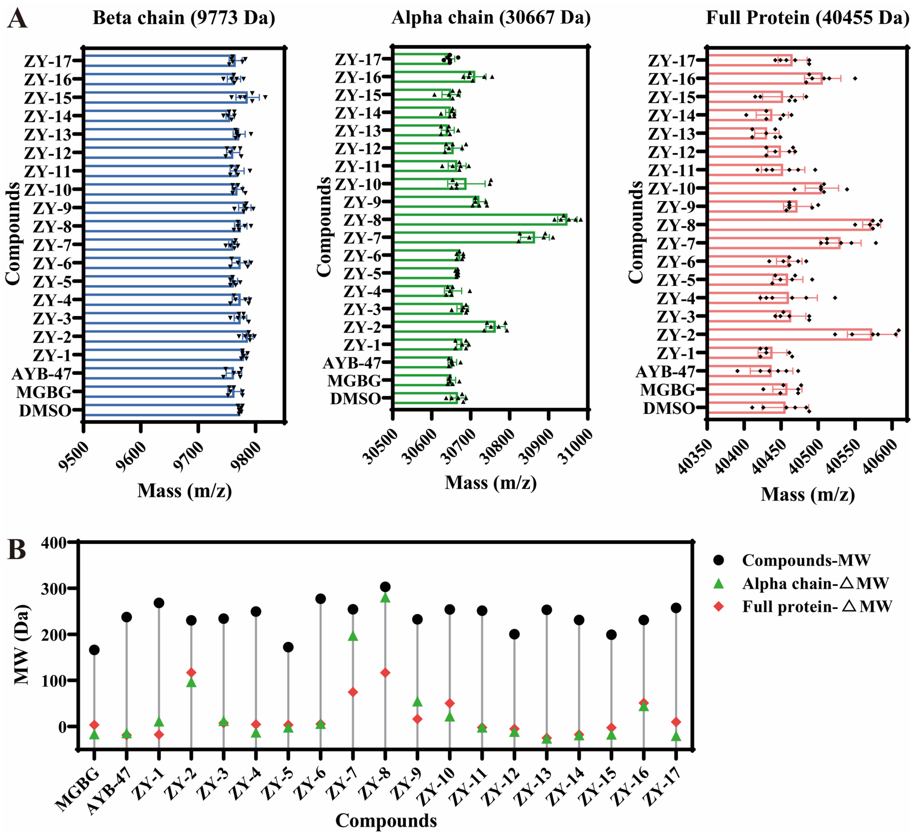


**Figure S7.** (A) The m/z values of the wild-type AdoMetDC protein determined by MALDI-TOF after being incubated (molar ratio 1:1) with the indicated compounds for 10 min. (B) The comparison of the molecular weights of the compounds and the mass changes of the wild-type AdoMetDC protein.

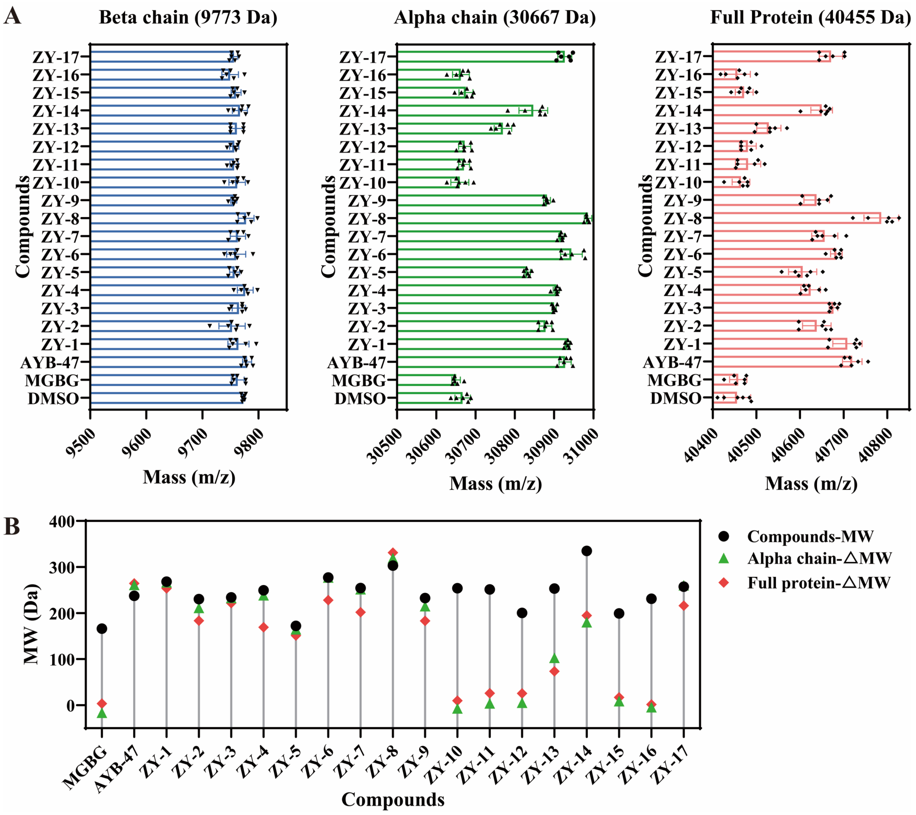


**Figure S8.** (A) The m/z values of the wild-type AdoMetDC protein determined by MALDI-TOF after being incubated (molar ratio 1:5) with the indicated compounds for 10 min. (B) The comparison of the molecular weights of the compounds and the mass changes of the wild-type AdoMetDC protein.

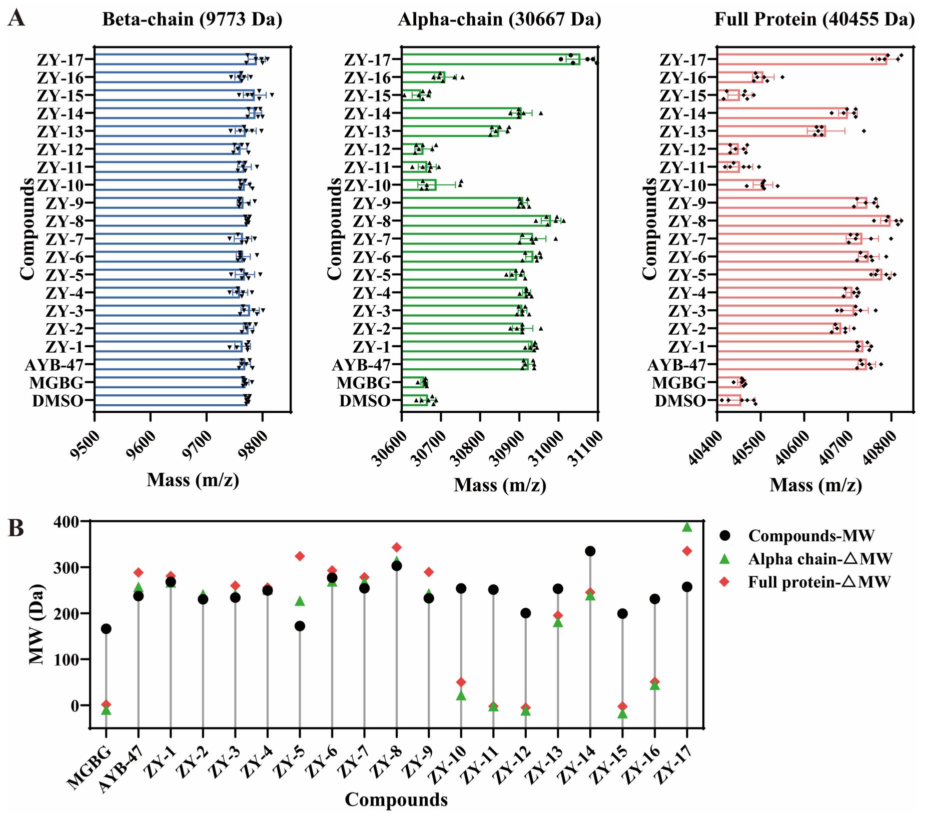


**Figure S9.** (A) The m/z values of the wild-type AdoMetDC protein determined by MALDI-TOF after being incubated (molar ratio 1:5) with the indicated compounds for 90 min. (B) The comparison of the molecular weights of the compounds and the mass changes of the wild-type AdoMetDC protein.

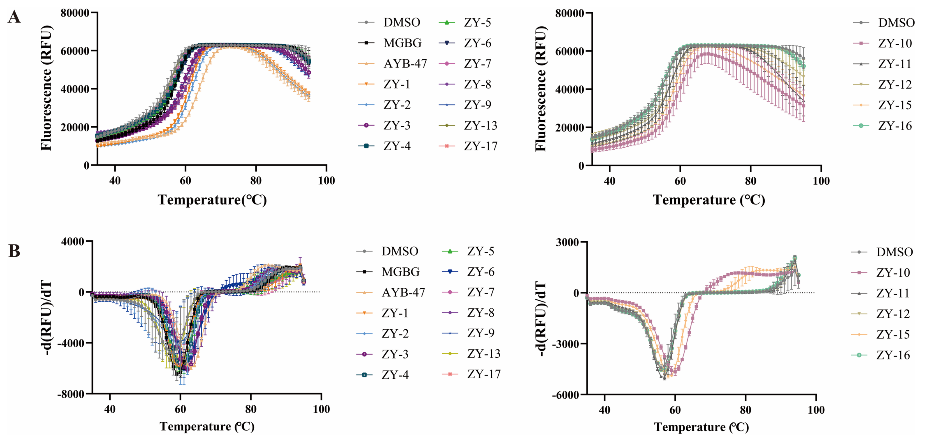


**Figure S10.** (A) The fluorescence intensity was recorded with the increase of the temperature in the presence of SYPRO Orange. (B) The data were transformed and plotted as a double-reciprocal Lineweaver-Burk plot (LB).

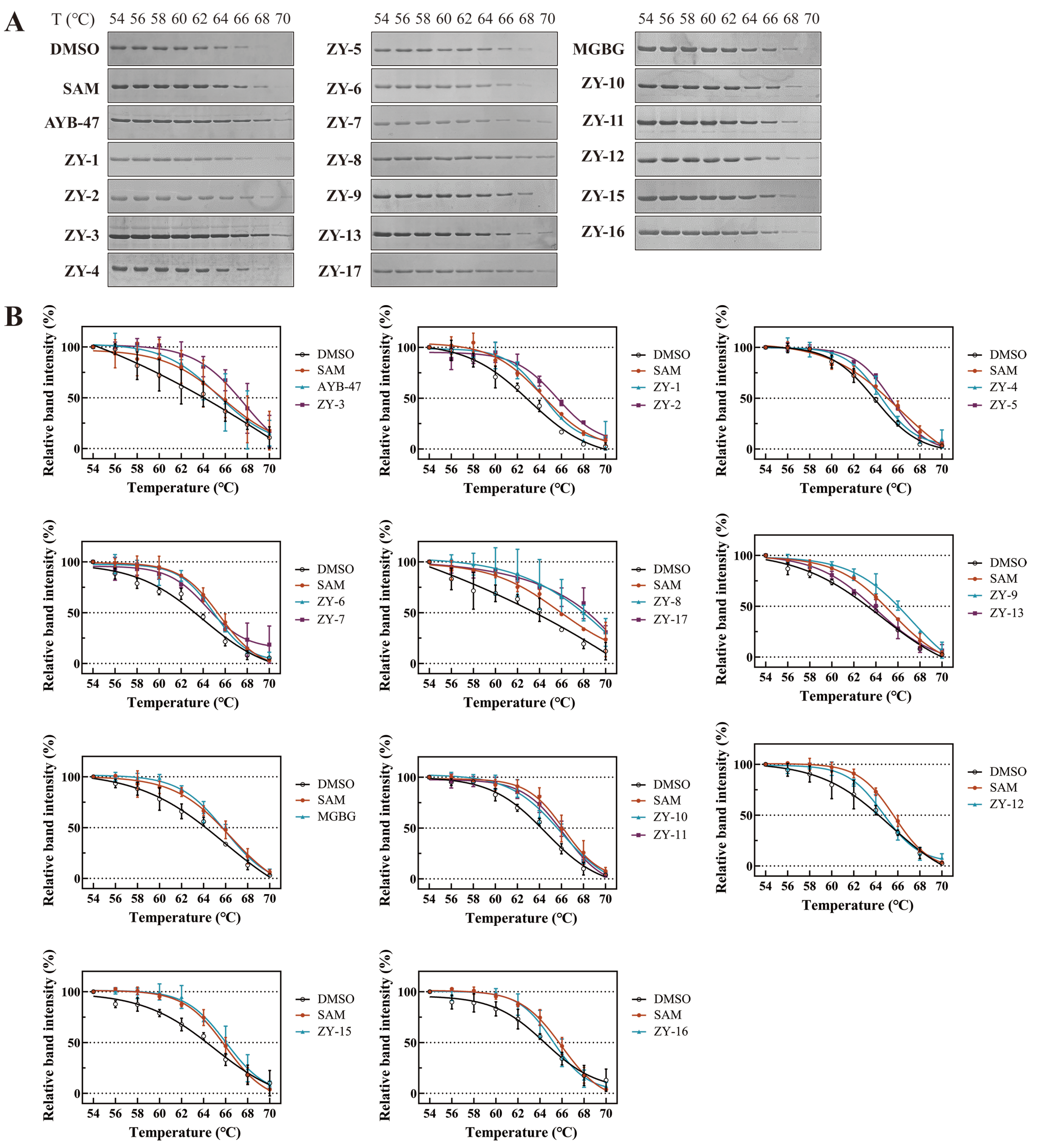


**Figure S11.** (A) Representative SDS-PAGE gels of the AdoMetDC protein samples treated with different inhibitors and temperatures. (B) The gray values of the AdoMetDC bands on the SDS-PAGE gels were calculated and melt curves for the inhibitors were generated using the Boltzmann S-type equation.

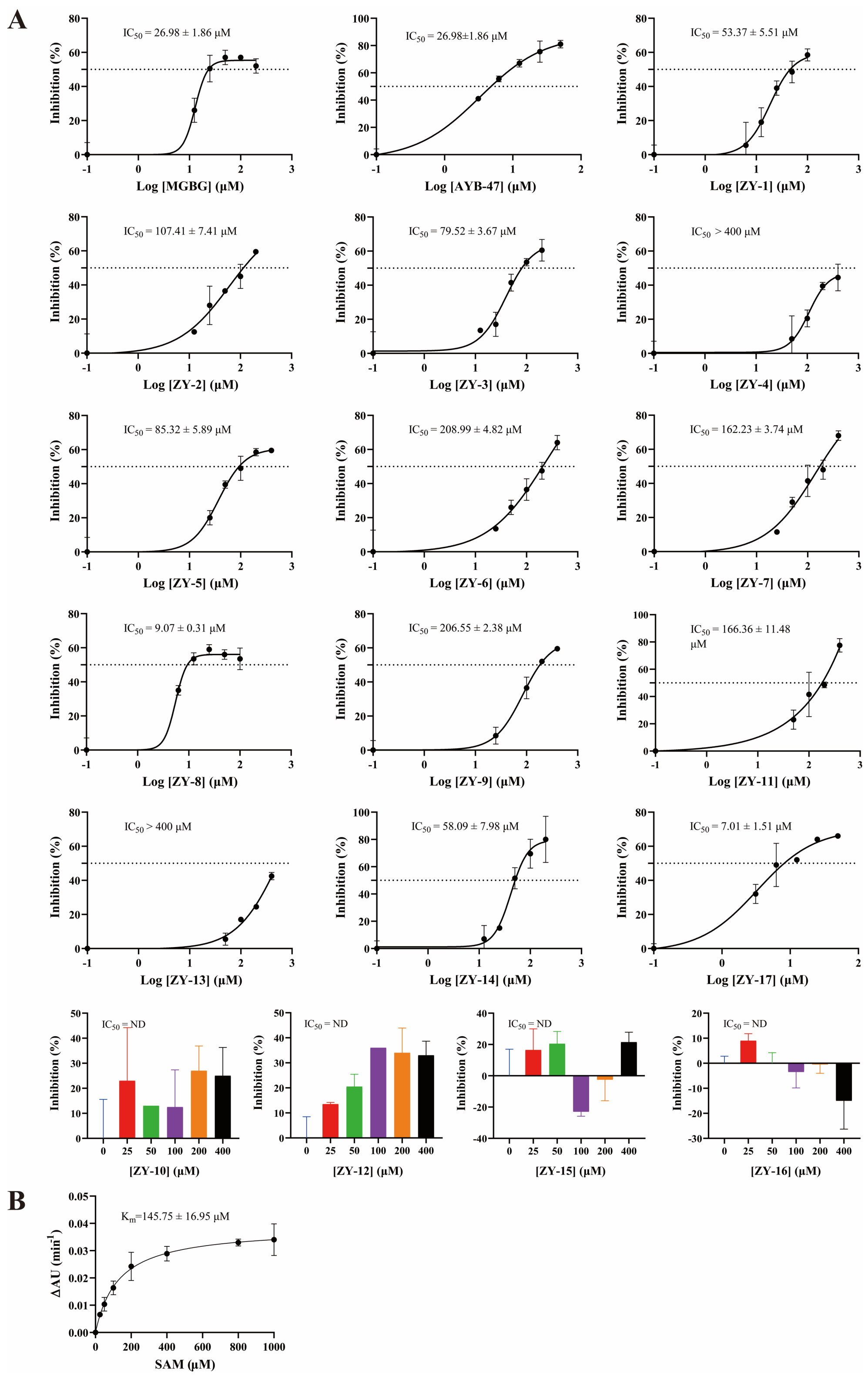


**Fig. S12** (A) The $IC_{50}$ values of the inhibitors were determined with the AdoMetDC-PEPC-MDH assay. Error bars represent standard deviations from two tests. (B) The $K_m$ value of the substrate AdoMet was determined by the AdoMetDC-PEPC-MDH assay.

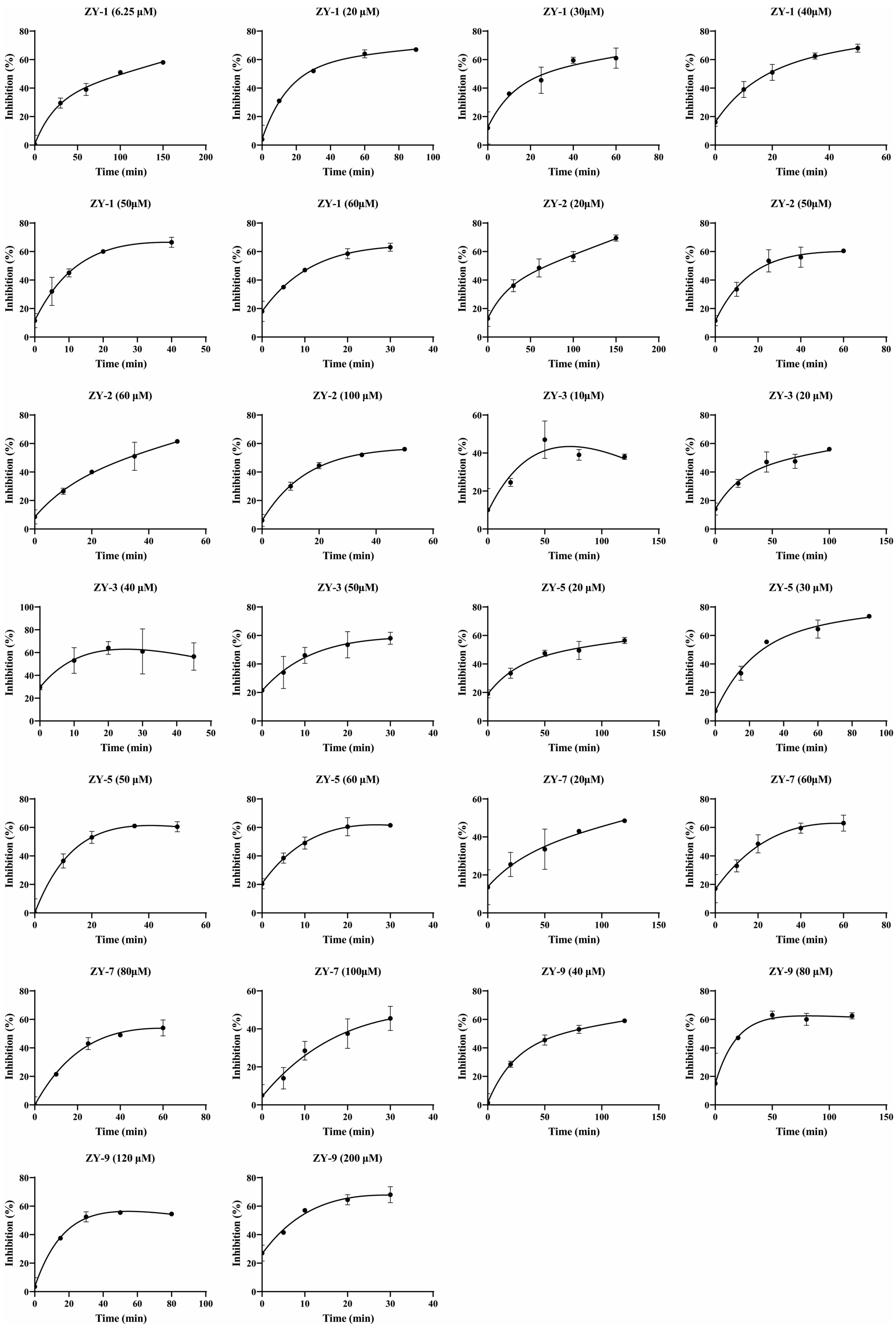


**Figure S13.** $k_{obs}$ values were determined by varying the pre-incubation time and the inhibitor concentration.

**Table S1**. The docking results and ionization potential data of known AdoMetDC inhibitors.

| PDB ID | Inhibitor name | Warhead group | Optimal pose score (kcal/mol) | Optimal pose ranking | Bonding atom distance (Å) | HOMO (Hartree) | HOMO (eV) | IP | N(HOMO) (%, SCPA) | N(HOMO), (eVx100, SCPA) | Local IP (x100) |
|---|---|---|---|---|---|---|---|---|---|---|---|
| **3DZ5** | M8M | $-CH_2-O-N^*H_2$ | -7.6 | 4 | 2.1 | -0.22501 | -6.1228 | 6.1228 | 1.67640 | -10.2643 | 10.2643 |
| **1I72** | MAO | $-CH_2-O-N^*H_2$ | -7.1 | 7 | 2.2 | -0.22448 | -6.1084 | 6.1084 | 1.29479 | -7.9091 | 7.9091 |
| **1I79** | MHZ | $-CH_2-NH-N^*H_2$ | -7.5 | 3 | 1.7 | -0.21743 | -5.9166 | 5.9166 | 0.16561 | -0.9798 | 0.9798 |
| **1I7B (3DZ3)** | SMM (MeAdoMet) | $-CH(N^*H_2)-CO-O-CH_3$ | -7.9 | 2 | 2.5 | -0.17443 | -4.7465 | 4.7465 | 0.05083 | -0.2413 | 0.2413 |
| **n.a.** | AbeAdo (MDL73811) | $-CH=CH-CH_2-N^*H_2$ | -7.4 | 9 | 2.2 | -0.22822 | -6.2102 | 6.2102 | 0.03372 | -0.2094 | 0.2094 |
| **n.a.** | **8** | $-CO-NH-N^*H_2$ | -8.6 | 3 | 3.4 | -0.23659 | -6.4379 | 6.4379 | 0.67516 | -4.3466 | 4.3466 |
| **n.a.** | mcule3265041117 | $-CO-NH-N^*H_2$ | -8.1 | 7 | 2.1 | -0.24260 | -6.6015 | 6.6015 | 0.17619 | -1.1631 | 1.1631 |
| **n.a.** | MCULE3626300541 | $-CO-NH-N^*H_2$ | -9.0 | 1 | 2.0 | -0.21048 | -5.7275 | 5.7275 | 0.51999 | -2.9782 | 2.9782 |
| **n.a.** | MCULE4717492978 | $-CO-NH-N^*H_2$ | -8.4 | 1 | 0.9 | -0.23986 | -6.5269 | 6.5269 | 0.14503 | -0.9466 | 0.9466 |
| **3DZ6** | M8E | $-CH_2-O-N^\#H_2$ | -7.3 | 3 | 1.9 | -0.22279 | -6.0624 | 6.0624 | 0.02022 | -0.1226 | 0.1226 |
| **1I7M** | CG (SAM486A) | $-NH-C(N^\#H_2)=NH$ | -9.0 | 3 | 3.0 | -0.22616 | -6.1541 | 6.1541 | 0.00765 | -0.0471 | 0.0471 |
| **3DZ4** | C8M | $-CH_2-CO-N^\#H_2$ | -7.8 | 3 | 2.6 | -0.22200 | -6.0409 | 6.0409 | 0.00481 | -0.0291 | 0.0291 |
| **3DZ2** | A8M | $-CH_2-N^\#H_2$ | -7.4 | 6 | 3.2 | -0.22355 | -6.0831 | 6.0831 | 0.00285 | -0.0173 | 0.0173 |
| **3DZ7** | O8M | $-CH_2-O-N^\#H_2$ | -6.9 | 12 | 4.0 | -0.23544 | -6.4067 | 6.4067 | 2.99948 | -19.2166 | 19.2166 |
| **3H0V** | M2T | N.A. | -6.6 | 8 | n.a | -0.18808 | -5.1179 | 5.1179 | N.A. | N.A. | N.A. |
| **3H0W** | N8M | N.A. | -7.7 | 3 | n.a | -0.22600 | -6.1498 | 6.1498 | N.A. | N.A. | N.A. |
| **1I7C** | MGB (MGBG) | $-NH-C(N^\#H_2)=NH$ | -6.6 | 3 | 2.8 | -0.21923 | -5.9656 | 5.9656 | 0.98886 | -5.8991 | 5.8991 |

Notes:

(1) All ligands were docked to the corresponding SCAR structures prepared from the original PDB structures. For ligands without PDB structures, they were docked to the SCAR structure of 3DZ5.

(2) The bonding atom distance was calculated as the bonding atom of the ligand and the bonding atom of the residue PYR68 in the original structure.

(3) The ligands were prepared with RDKit for 3D conformation generation.

(4) M2T and N8M do not contain primary amines for covalent bonding.

(5) Optimal pose: The first docked conformations with the scaffold structure fitting with the ligand structure in the crystal structure.

(6) Docking results were from our previous data [1]. A difference is in the ranks of the optimal poses. In our previous paper, poses with same docking scores were given same ranks. In this table, the ranks are the model numbers from the AutoDock Vina outputs.

(7) *: The bonding atom in the covalent inhibitor. #: The atom used for calculating the partial HOMO value in non-covalent inhibitors.

(8) N.A.: not available.

(9) IP: ionization potential.

**Table S2**. The docking results and ionization potential data of 18 compounds purchased for experimental validation. The last one was not tested because of low solubility.

| Screening ID | Optimal Docking Score | Rank | Bonding atom distance (Å) | HOMO (Hartree) | HOMO (eV) | IP | N(HOMO) (%, SCPA) | N(HOMO), (eVx100, SCPA) | Local IP (x100) |
|---|---|---|---|---|---|---|---|---|---|
| **ZY-1** | -8.5 | 7 | 1.1 | -0.2491 | -6.7775 | 6.7775 | 0.0973 | -0.6597 | 0.6597 |
| **ZY-2** | -8 | 3 | 1.0 | -0.2734 | -7.4396 | 7.4396 | 8.5233 | -63.4097 | 63.4097 |
| **ZY-3** | -8.2 | 1 | 2.2 | -0.2691 | -7.3218 | 7.3218 | 0.6213 | -4.5489 | 4.5489 |
| **ZY-4** | -9.1 | 5 | 1.9 | -0.2305 | -6.2711 | 6.2711 | 0.0681 | -0.4269 | 0.4269 |
| **ZY-5** | -8.1 | 1 | 2.1 | -0.2332 | -6.3449 | 6.3449 | 0.3805 | -2.4143 | 2.4143 |
| **ZY-6** | -8 | 1 | 2.1 | -0.2503 | -6.8121 | 6.8121 | 0.0087 | -0.0589 | 0.0589 |
| **ZY-7** | -8.1 | 9 | 1.0 | -0.2140 | -5.8243 | 5.8243 | 0.1069 | -0.6224 | 0.6224 |
| **ZY-8** | -8.3 | 7 | 0.9 | -0.2416 | -6.5735 | 6.5735 | 0.0152 | -0.1000 | 0.1000 |
| **ZY-9** | -8.2 | 3 | 2.2 | -0.2527 | -6.8769 | 6.8769 | 0.0803 | -0.5524 | 0.5524 |
| **ZY-10** | -8 | 4 | 1.0 | -0.2570 | -6.9941 | 6.9941 | 0.1139 | -0.7964 | 0.7964 |
| **ZY-11** | -8.8 | 3 | 2.0 | -0.2629 | -7.1533 | 7.1533 | 1.0432 | -7.4626 | 7.4626 |
| **ZY-12** | -7.6 | 6 | 1.8 | -0.2113 | -5.7492 | 5.7492 | 0.5041 | -2.8979 | 2.8979 |
| **ZY-13** | -8.1 | 3 | 1.6 | -0.2699 | -7.3438 | 7.3438 | 1.5356 | -11.2775 | 11.2775 |
| **ZY-14** | -8.5 | 5 | 1.5 | -0.2388 | -6.4981 | 6.4981 | 1.4207 | -9.2318 | 9.2318 |
| **ZY-15** | -8.1 | 1 | 1.9 | -0.23096 | -6.2847 | 6.2847 | 0.03588 | -0.2255 | 0.2255 |
| **ZY-16** | -8.1 | 8 | 0.9 | -0.23259 | -6.3291 | 6.3291 | 1.38584 | -8.7711 | 8.7711 |
| **ZY-17** | -10.2 | 1 | 1.1 | -0.2038 | -5.5460 | 5.5460 | 0.0340 | -0.1886 | 0.1886 |
| / | -8.3 | 5 | 2.0 | -0.22143 | -6.0254 | 6.0254 | 6.38159 | -38.4518 | 38.4518 |
